# Effect of surfactants on SARS-CoV-2: Molecular Dynamics Simulations

**DOI:** 10.1101/2022.11.17.516905

**Authors:** Marc Domingo, Jordi Faraudo

## Abstract

Surfactants are commonly used as disinfection agents in personal care products against bacteria and viruses, including SARS-CoV-2. However, there is a lack of understanding of the molecular mechanisms of the inactivation of viruses by surfactants. Here, we employ coarse grain (CG) and all-atom (AA) molecular dynamics simulations to investigate the interaction between general families of surfactants and the SARS-CoV-2 virus. To this end, we considered a CG model of a full virion. Overall, we found that surfactants have only a small impact over the virus envelope, being inserted into the envelope without dissolving it or generating pores, at the conditions considered here. However, we found that surfactants may induce a deep impact on the spike protein of the virus (responsible for its infectivity), easily covering it and inducing its collapse over the envelope surface of the virus. AA simulations confirmed that both negatively and positively charged surfactants are able to extensively adsorb over the spike protein and get inserted into the virus envelope. Our results suggest that the best strategy for the design of surfactants as virucidal agents will be to focus on those strongly interacting with the spike protein.

## I. INTRODUCTION

The SARS-CoV-2 coronavirus emerged in December 2019 as a human pathogen which causes the COVID-19 respiratory disease^1,2^ and at the time of writing this manuscript, the virus is still with us. Furthermore, COVID-19 is not the first pandemic that has been caused by a human coronavirus with zoonotic origin, being the previous cases SARS-CoV (2003) and MERS (2012) and future spillovers seem likely^3^. Therefore, investigations on basic scientific questions related to coronavirus inactivation and disinfection are of great importance. Here we will consider a computational study with the aim to provide rational tools to select surfactants for disinfection processes suitable for SARS-CoV-2.

Coronaviruses are enveloped virus, which means that the genetic material of the virus is protected by an envelope. In the case of SARS-CoV-2, the envelope is formed by phospholipids, membrane (M) proteins and envelope (E) proteins. The envelope also has glycosilated spike (S) proteins that protrude from the envelope and are responsible for the infectivity of the virus.

The transmission of respiratory viruses like SARS-CoV-2 involves the expiration to the medium by an infected individual of droplets or aerosols that contain virus particles. The infection of another individual may be divided into two different mechanisms, direct or indirect transmission. The direct mechanism involves inhalation of aerosols or the deposition of emitted droplets on mucosal surfaces. The indirect mechanism means that an expiratory droplet containing virus may land into an environmental surface. Coronaviruses can remain viable over surfaces for extended periods of time (hours or days depending on the material) and eventually another individual may touch the contaminated surface and consequently get infected when touching his/her mucoses^3–5^.

In the case of SARS-CoV-2, the studies developed during the present pandemic show that it is able to remain infectious over many different surfaces and remains stable under a wide range of environmental conditions^6,7^. For example, it remains infectious for 72h over plastic surfaces^5^, 7 days at a surgical mask^6^ and between 9h to 22h over human skin depending on the virus variant^8,9^. Our MD simulations indicate that the virus spikes can be attached to sebaceous human skin without any deformation or alteration and retain their hydration^10^. Also, our simulations indicate that they can remain adsorbed with a high affinity over common plastics without any substantial deformation^11^. Only some materials such as metals^12^ or carbon based materials^13^ seem to be able to have a substantial impact on the adsorbed virus spikes. These evidences justify the recommendations by health agencies about cleaning and disinfection of surfaces and hands^14^.

One of the most typical disinfecting agents are surfactants which are employed in the formulation of soaps and other household cleaning products. Many common surfactants (such as cationic quaternary ammonium surfactants or anioninc sulfate surfactants) are effective against different pathogens including viruses^15–18^. In the case of SARS-CoV-2, experimental evidence showed inactivation by Sodium Laureth Sulfate, typically employed in soap^18^. Interestingly, experiments done using commercial hand soap indicate that SARS-CoV-2 is able to remain infectious after 5 min exposure to hand soap but not after 15 min exposure^6^.

Little is known about the mechanisms and interactions that occur when inactivating viruses by means of surfactants^19^, precluding its rational design for disinfecting applications. It is known that surfactants are able to denaturize or solubilize proteins and also they are able to destabilize lipid membranes due to its hydrophobic/hydrophilic nature^20–23^. However, it is not clear how these effects may contribute to inactivate an enveloped virus such as SARS-CoV-2. In principle, we can suggest as possible mechanisms the denaturation of the spike protein or the damage of the integrity of the virus envelope (Figure 1) but further research is needed in order to support or disprove them.

**FIG. 1.**
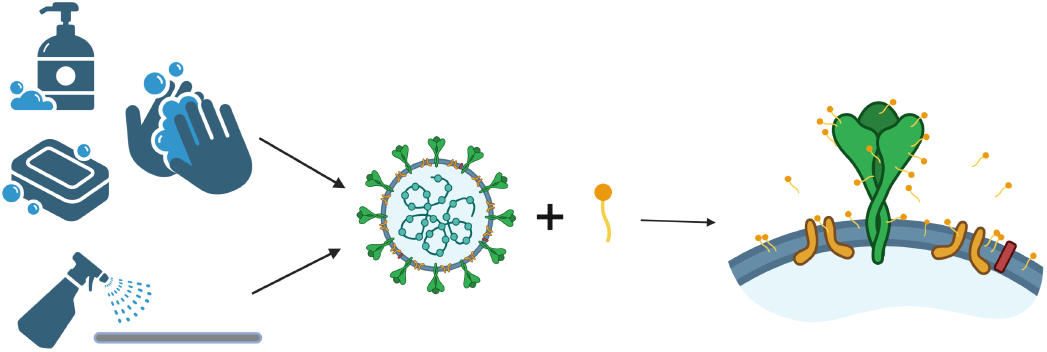
Scheme of the possible interaction between SARS-CoV-2 virus and surfactants. Made with BioRender.com.

In order to investigate the possible interactions between surfactants and SARS-CoV-2 virus from a fundamental physico-chemical point of view, we perform here molecular dynamics simulations of this system. First, we use a coarse-grain (CG) model to perform molecular dynamics simulations of a SARS-CoV-2 virion in the presence of different surfactants. After that, we consider simulations with full atomic resolution. In this case, we consider a patch of a virus envelope membrane made of lipids and a full spike protein in contact with different surfactants. Overall, our simulations indicate that the most relevant mechanism in the interaction between surfactants and the virus is the interaction with the spike protein. Also, our atomistic simulations suggest a higher interaction of the spike with anionic surfactants as compared with cationic surfactants.

## II. METHODS

### A. Coarse-Grain (CG) Molecular Dynamics Simulations

#### 1. CG Models for the virus and surfactants

We have performed Coarse-Grained (CG) MD simulations of a full SARS-CoV-2 virus in the presence of surfactants using LAMMPS 19 Sep 2019 version^24^. Our simulations employ a CG model of surfactants developed by^25^ and a compatible CG model of a full virion particle developed by^26^ (see Figure 2). All the elements of the employed CG models of the virus and the surfactants and their interactions are summarized schematically in Figure 3 and described in detail below.

**FIG. 2.**
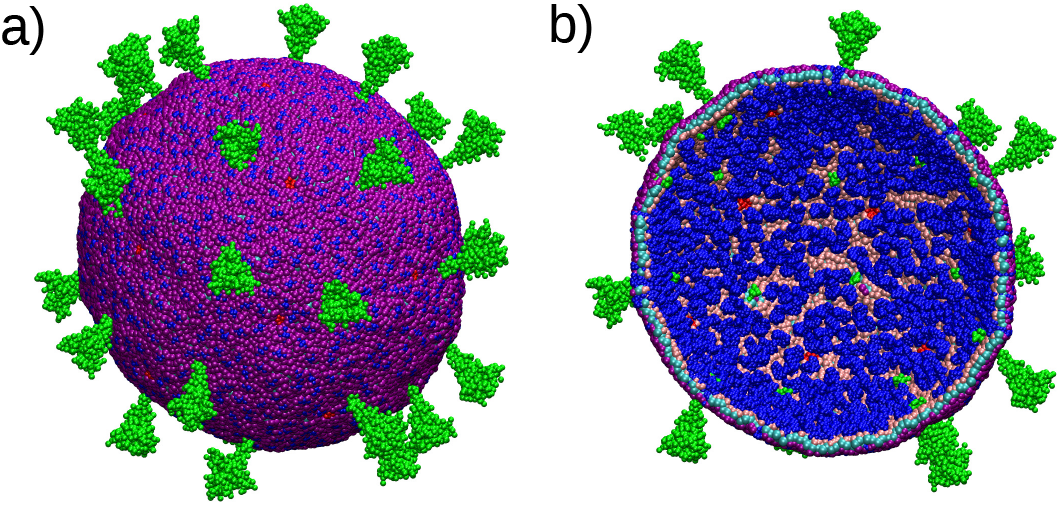
Coarse grain model of the SARS-CoV-2 envelope developed by^26^. Spike protein in green, M protein in dark blue, E protein in red, viral lipids in purple, cyan and pink. a) View of the full virus particle; b) Cut of the view in a) so that the internal structure of the envelope can be seen.

**FIG. 3.**
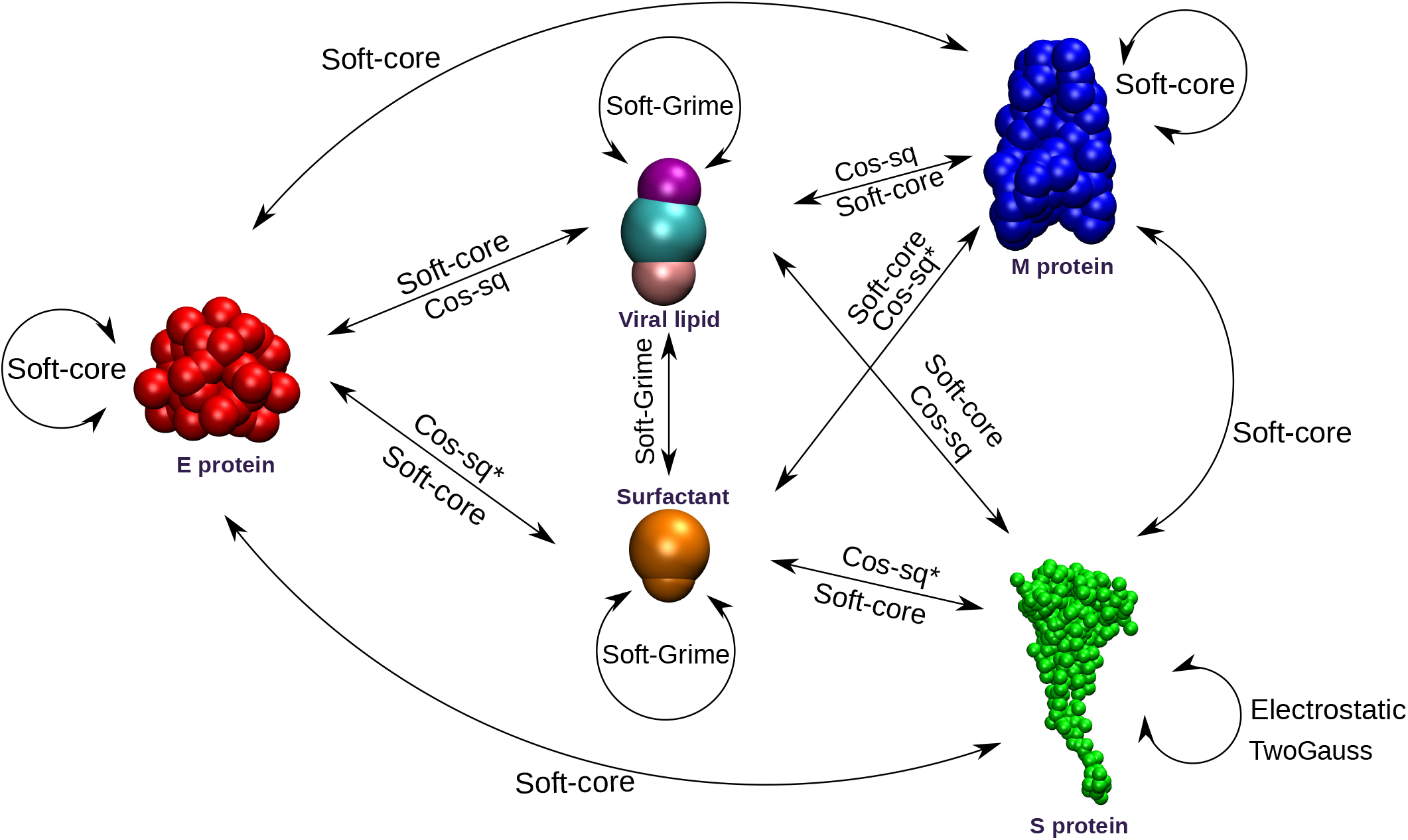
Scheme of the non-bonded interactions between the components of the virus and surfactants used for the simulations. (*) Interaction not considered for simulations 1, 2 and 3 in table I. Surfactant in orange color.

The CG model of the SARS-CoV-2 virus considers the virus envelope (Figure 2) and it neglects the internal structure of the virus (the viral RNA and the N proteins that encapsulate the genetic material inside the virion are not included in the model). In this model, the virus envelope is made of 1000 dimeric membrane (M) proteins and 20 envelope (E) pentameric transmembrane proteins in a lipid bilayer of 38580 lipids. It has also 30 Spike trimeric proteins (S) inserted at the envelope by a small transmembrane fragment (see Figure 2).

The virion proteins M and E were coarse-grained in a way that each CG bead corresponds approximately to 5 aminoacids. All CG beads of M and E protein have zero charge. The structure of these proteins is maintained by rigid bonds. The interactions between beads of different proteins were modelled using the following soft core cosine potential that avoids overlap between beads:

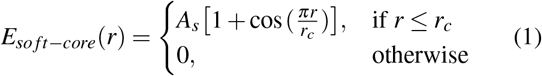

where *r_c_* is equal to the bead size. We take the same values of *A_s_* and *r_c_* as in^26^. In the case of the S protein, the S1 and S2 domains were mapped to 60 and 50 CG beads respectively, and the 22 glycans of S were each mapped to a single bead. Some beads of S are charged. Interactions between beads of the same S protein (intraprotein interactions) were modelled using a heteroelastic network model. Interprotein interactions were composed of excluded volume, attractive, and screened electrostatic terms. Excluded volume interactions were modelled using the soft core interaction given by Eq.(1) as for M and E proteins. Attractive, nonbonded interactions between interprotein contacts were modeled as the sum of two Gaussian potentials:

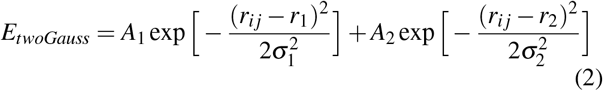

where the parameters *r*_1_, *r*_2_, σ_1_ and σ_2_ were obtained from all atomic simulations (see^26^). Screened electrostatic interactions were modeled using Yukawa potentials,

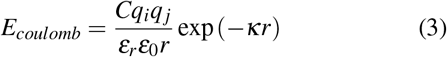

where *q_i_* is the charge of CG bead *i*, *κ* = 1.274 nm^-1^ is the inverse Debye length for 150 mM of monovalent salt, and *ε_r_* = 17.5 is the effective dielectric constant of the protein environment.

The lipids of the virus envelope were modelled using the bilayer CG model previously developed in^25^ as done by^26^. This bilayer model consists of 3 beads per bilayer segment (i.e. “1.5” beads/lipid), with two hydrophilic headgroup beads (heads) connected by harmonic bonds to a hydrophobic bead designed as “interfacial bead” in^25^ (see Figure 3). The interaction potential between lipid beads has a soft core repulsion and a short range attraction:

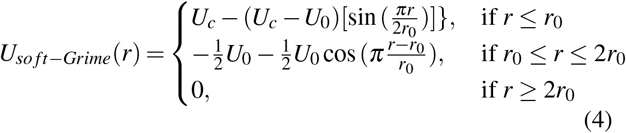

In Eq.(4), r0 is the size of the bead, *U_c_* is a large repulsive energy preventing bead overlap and −*U*_0_ is the depth of the energy minima at *r* = *r*_0_. We note here that since Eq.(4) is not implemented in the standard release of LAMMPS, we have employed the custom code available in Ref^27^ (see details in the Appendix).

In the virus model employed in the simulations (Figure 2) we have 19290 interfacial beads and 38580 head beads, modelling 38580 fosfolipids. The size of the beads was selected to be *R_I_* = 1.8 nm for the interfacial beads and *R_H_* = 0.75R_*I*_ for the head beads^26^. The characteristic energies *U_c_* and *U*_0_ are the same as in^26^ and depend on the type of bead. For the interaction between interfacial beads we have:

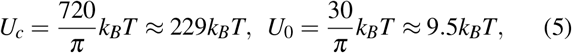

where the *π* factors are introduced for convenience in the calculation of the force (see Appendix). The interaction between head beads is considered purely repulsive:

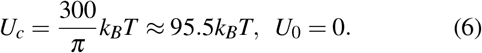

The cross interaction between head and interfacial beads was also considered to be purely repulsive and given by Eq.(5) with *r*_0_ = *R_I_* = 1.8 nm. Using these values, the model lipids self-assemble spontaneously into a vesicle^25^.

Cross interactions between the lipids and the structural proteins M,E and S are responsible for the stability of these proteins in the envelope. These interactions were modelled using an attractive potential between lipid beads and the transmembrane domains of M, E and S proteins given by:

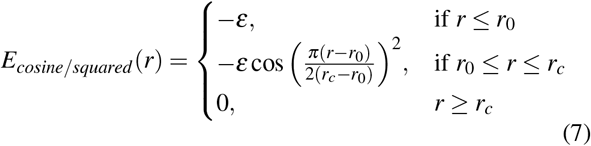

In Eq.(7), *r*_0_ is the sum of the radii of the beads that interact and *r_c_* is the cutoff which is set *r_c_* = 2*r*_0_. Unless stated otherwise, we will consider *ε* = 4 kcal/mol= 6.7*k_B_T* as in Ref^26^. In order to explore the effect of a different strength of the lipid-protein interaction we will also consider additional simulations with a weaker interaction with *ε*=0.63 kcal/mol = 1.06*k_B_T* as considered in the original paper introducing this interaction^28^. We checked that for both values of ε the structure of the virus model shown in Figure 2 remains stable (proteins remain inserted in the bilayer envelope).

In order to model the surfactants that will interact with the virus in a way consistent with the virus CG model, we used also the CG model of^25^ for describing the surfactants (see Figure 3). This model was originally proposed as a generic and flexible CG model for describing self-assembly of amphiphilic molecules in vesicles, micelles and other structures and includes both lipids and surfactants. In this model, surfactants always have zero charge since electrostatic interactions were not considered explicitly in the model^25^.

Each CG surfactant has two beads, one with hydrophilic character (head bead) and another with hydrophobic character (tail bead). The size of the surfactant tail beads is taken equal to the size of the lipid interfacial beads, *R_I_* = 1.8 nm. The size of the surfactant head beads is taken as *R_SH_* = 1.5*R_I_* because for this combination of bead sizes, the CG surfactants form micelles^25^. The interaction between surfactants was described, as in the case of lipids, by Eq.(4), using the same parameters as in the lipid case (Eqs. (5)–(6)).

The surfactant - lipid cross interaction (figure 3) was described also by the soft potential given by Eq. (4). As in the case of the lipid-lipid and surfactant-surfactant interactions, the interactions of head beads with any other surfactant or lipid bead is purely repulsive. In order to model different types of surfactants (different strengths for the lipid-surfactant interaction) we have considered several different *U*_0_ strengths for the attractive depth (Eq.(4)) in the interactions between surfactant beads of tail type and lipid beads of interfacial type. We have also adjusted the value of *U_c_* for the different values of *U*_0_ in Eq.((4) to avoid overlap between beads for the cases with strongest attraction. The surfactant - protein interaction was modelled analogously to the lipid - protein interaction using Eq. (7) for all protein-surfactant beads, exploring different values of ε to mimic different degrees of interaction between surfactants and proteins.

#### 2. Description of CG simulations

The simulated system consists of a SARS-CoV-2 virus particle and 4000 surfactant molecules in a simulation box of dimensions 240×240×240 nm^3^, which corresponds to a surfactant concentration of 0.5 mM. In all the simulations, we have integrated the equations of motion using a timestep of 100 fs as in^26^. The temperature of the system was set at 300K. We employed a Langevin thermostat as implemented in LAMMPS^24,29^ with a decay time of *t_damp_* = 100 ps as in^25^.

We have performed simulations exploring different values of the strength of the interactions between surfactants and the virus components (proteins and lipids), as indicated in Table I (Simulations 1 to 9). As seen in the table, the surfactant-lipid interactions ranged from an attraction equal to that of the lipid-lipid interaction to a much larger attraction and the surfactant protein interactions ranged from no interaction at all to an attraction equal to that of the lipid-protein interaction.

**TABLE I.** Force Field parameters in CG simulations. We indicate the values of the parameters for the interactions between surfactants and lipids (*U*_0_ in Eq.(4)), surfactants and proteins (*ε* in Eq. 7) and lipids and proteins (*ε* in Eq. 7).

| Sim. | surf-lipid<br>$U_0/k_B T$ | surf-protein<br>$\epsilon/k_B T$ | lipid-protein<br>$\epsilon/k_B T$ | Comments |
| --- | --- | --- | --- | --- |
| 1 | 9.5 | 0 | 6.7 | No surfactant-protein interaction.<br>(surfactants only interact with lipids) |
| 2 | 28.5 | 0 | 6.7 |  |
| 3 | 95.0 | 0 | 6.7 |  |
| 4 | 9.5 | 1.06 | 6.7 | Weak surfactant-protein interaction |
| 5 | 28.5 | 1.06 | 6.7 |  |
| 6 | 95.0 | 1.06 | 6.7 |  |
| 7 | 9.5 | 6.7 | 6.7 | Surfactant-protein interaction equal<br>to lipid-protein interaction |
| 8 | 28.5 | 6.7 | 6.7 |  |
| 9 | 95.0 | 6.7 | 6.7 |  |
| 7' | 9.5 | 1.06 | 1.06 | Modified SARS-CoV-2 model<br>(weaker lipid-protein interaction)) |
| 8' | 28.5 | 1.06 | 1.06 |  |
| 9' | 95.0 | 1.06 | 1.06 |  |

In order to test the influence of the parameters employed for simulating the supramolecular structure of the SARS-CoV-2 virus (in particular, the strength of the lipid-lipid interaction) we also ran a series of three additional simulations in which we weakened the lipid-protein interaction. In these simulations, the surfactant-protein interaction was considered equal tot he lipid-protein interaction. These simulations are indicated as 7’, 8’ and 9’ in Table I.

The initial configuration of the system was generating employing the equilibrated virus structure provided in^26^. The surfactants were added using LAMMPS scripting options.

The simulation time in all cases was selected so that key physical quantities of interest (potential energy, number of adsorbed surfactants,…) are equilibrated. The required simulation times were between 1.3×10^7^ - 3.0×10^7^ time steps.

All analysis and images were done using VMD^30^.

#### 3. Replica exchange (RE) simulation

Convergence to metastable states is always an issue in MD simulations (particularly in CG models). In order to test possible lack of convergence to equilibrium of our simulations, we did a test using the Replica Exchange (RE) method. Starting with the same initial condition and parameters as in Simulation 7 in Table I, we have performed a Replica Exchange (RE) MD simulation in which we considered 24 different replicas separated by 20K each one (*T* of the thermostats ranged from 300K to 760K), with a frequency trial of a swap between replicas of each 100 time steps. This simulation was run for 1 · 10^6^ time steps.

The result from the RE simulation was used as starting point for an unbiased, standard MD simulation at 300K. From the 24 final configurations obtained in the RE simulation, we took the one corresponding to the replica at 300K as initial configuration for the new simulation. This additional MD simulation at 300K after the RE lasted for 1.5 · 10^7^ and the interaction parameters used are also the ones from simulation 7 in Table I.

### B. All Atomic (AA) simulations

We have performed all atomic MD simulations of the SARS-CoV-2 Spike glycoprotein inserted into a bilayer membrane (corresponding to a small patch of the virus envelope) in contact with two different surfactants, the cationic Dodecyltrimethylammonium (DTAB) and the anionic Sodium Dodecyl Sulfate (SDS).

The structure of the fully glycosilated and palmitoylated S protein (“up” conformation) inserted into a membrane was taken from a previously developed model^31^ and it is shown in Figure 4a). The lipid membrane were the S protein is inserted is constituted by a mix of POPC, POPE, POPS, POPI and cholesterol lipids mimicking the composition of the endoplasmic reticulum-Golgi intermediate compartment^31^. The chemical structures of the surfactants considered in our simulations were shown in Figure 4b and Figure 4c.

**FIG. 4.**
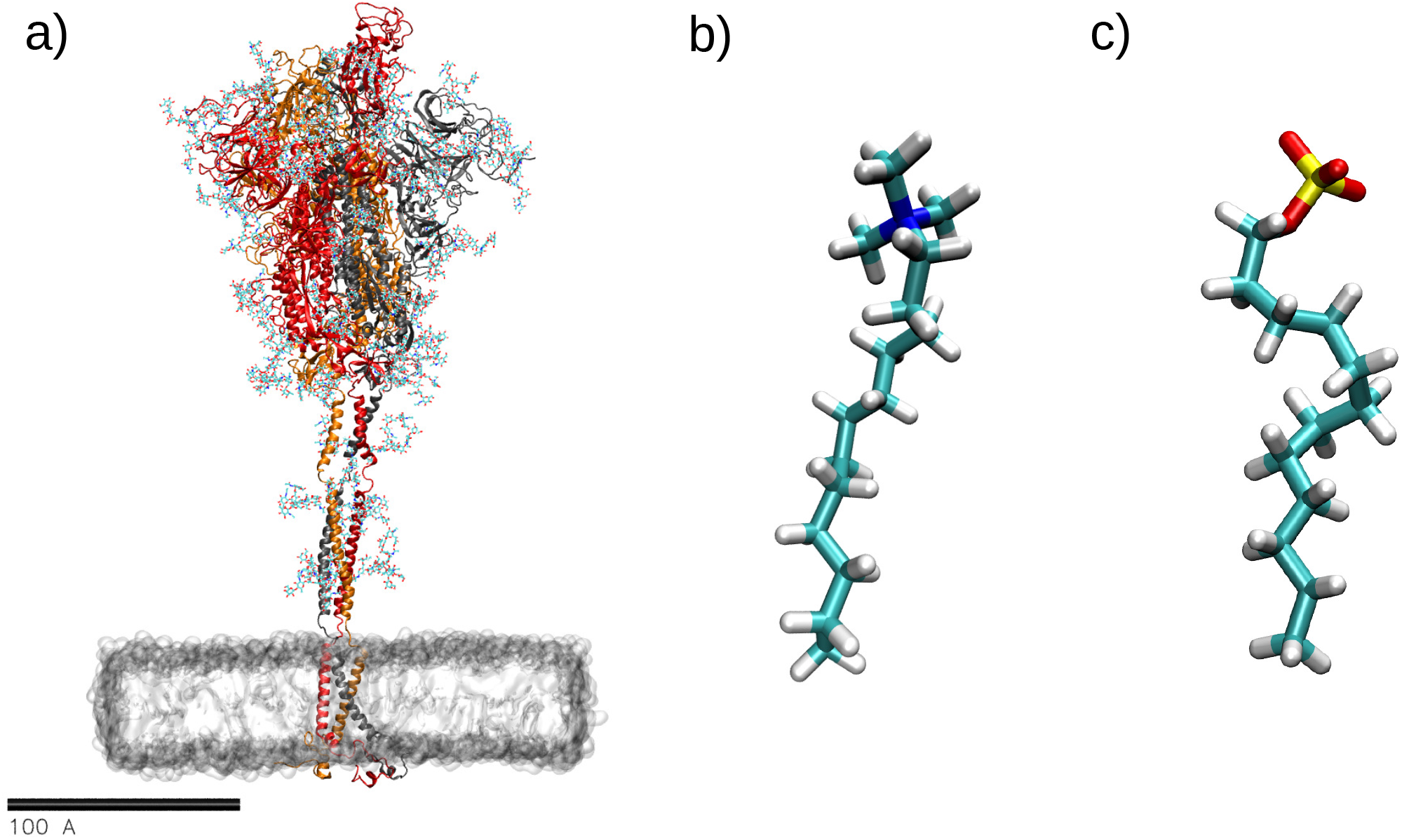
Structures used in AA simulations. a) Full spike glycoprotein inserted into the membrane. Model from^31^. Protein in NewCartoon representation (monomers in red, orange and gray color), glycans in line representation and membrane represented by a surface; b) Dode-cyltrimethylammonium bromide surfactant (DTAB); c) Sodium dodecyl sulfate surfactant (SDS). Atom colors: Carbon in cyan, hydrogen in white, nitrogen in blue, oxygen in red, sulfur in yellow.

The MD simulations were performed using NAMD 2.14 software^32^. the equations of motion were integrated with a 2 fs time step and electrostatic interactions were updated every 4 fs. All bonds between heavy atoms and hydrogen atoms were kept rigid. In all simulations we employed periodic boundary conditions in all directions. Lennard-Jones interactions were computed with a cutoff of 1.2 nm and switching function starting at 1.0 nm. Electrostatic interactions were computed using Particle Mesh Ewald (PME) algorithm using a real space cutoff set at 1.2 nm and a PME grid at 1.0 Å. The temperature was set at 300K in all simulations, employing the Langevin thermostat with a damping coefficient of 1 ps^-1^. In NPT simulations we employed a Nosé-Hoover isotropic barostat with an oscillation period of 100fs and a damping time of 50fs.

The force-field employed in the simulations was CHARMM36^33,34^ as in^31^ and as in our previous simulations of the SARS-CoV-2 Spike glycoprotein^10,11,13^. The force field also includes an appropiate parametrization for the surfactants.

The initial coordinates for the simulations where obtained as follows. The structure of S inserted into a membrane was downloaded from^31^. SDS and DTAB structures were prepared using CHARMM-GUI^35–37^ and distributed randomly in the water phase but only in the external side of the membrane exposing the S protein. The box was solvated with water molecules using VMD and neutralized with NaCl. Also, we added 150 mM of NaCl. A short NPT simulation for both systems was performed keeping the membrane surface constant (only the direction perpendicular to the membrane was allowed to pressurize), until we achieve the correct water density.

Once the system was equilibrated, we performed production NVT simulations. In order to avoid crossing of surfactants from the external side of the membrane to the internal side across the periodic boundary conditions, we added an empty space at both sides, generating water/vacuum interfaces that preclude the crossing of the surfactants. The details of the simulations are summarized in Table III.

**TABLE II.** Results from Simulations 1-9. We indicate the number of surfactants that remain free in solution and not in contact with any bead of the virus (we considered a cutoff of 7.5 nm between the surfactants and any virion bead to consider that the surfactant was free), the number of surfactants in contact with lipids and the number of surfactants in contact with Spike proteins at the final equilibrium state. In all simulations the total number of surfactants is 4000 (see Methods).

| Sim. | Free surfactants | In contact w/lipids | In contact w/S protein |
| --- | --- | --- | --- |
| 1 | 3793 | 206 | 0 |
| 2 | 3691 | 292 | 0 |
| 3 | 3682 | 305 | 0 |
| 4 | 0 | 708 | 1708 |
| 5 | 0 | 1565 | 1498 |
| 6 | 0 | 1830 | 1405 |
| 7 | 0 | 809 | 2502 |
| 8 | 28 | 1897 | 2340 |
| 9 | 90 | 2543 | 1793 |

**TABLE III.** Characteristic parameters of the AA simulations of the spike protein with surfactants.

| Simulated system | Num. surfactants | Box size (nm<sup>3</sup>) | Atoms | Na<sup>+</sup> | Cl<sup>-</sup> | Sim time (ns) |
| --- | --- | --- | --- | --- | --- | --- |
| Spike protein with DTAB | 110 | 20.47 × 19.94 × 50.85 | 1592846 | 1453 | 1282 | 188 |
| Spike protein with SDS | 128 | 20.47 × 19.94 × 50.85 | 1590529 | 1692 | 1283 | 127 |

During the simulations, we monitored the number of surfactants adsorbed over the S protein and also the number of surfactants inserted into the membrane. For both simulations a contact between surfactants and the S protein was considered if at least one atom of the surfactant molecule was within 3 Å of an atom constituting the spike protein, including the glycans. In order to count the number of surfactants in contact with the S protein at each time step, we employed a TCL script running on VMD^30^ implementing the distance requirement described above. In order to calculate the number of inserted surfactants into the membrane, the following approach was considered. First, we aligned the MD trajectory by considering the phosphorous atoms from the upper layer of the membrane. With the aligned trajectory, then we calculated the center of mass of the same set of phosphorous atoms (for each time step) and made the average. As the membrane is aligned perpendicular to the *z* axis, we considered that a surfactant was inserted into it if the *z* coordinate of the carbon at the extreme of the surfactant was lower than the z coordinate of the center of mass of the phosphorous atoms at the upper layer.

All images were generated using VMD^30^.

## III. RESULTS AND DISCUSSION

### A. CG simulation of the interaction of surfactants with a SARS-CoV-2 virion

To illustrate the results of our CG simulations, we first describe in detail a specific particular case (simulation 4 in Table I) and after that we will discuss how the results change for the different combinations of force field parameters considered in Table I.

In simulation 4 we have performed two different runs corresponding to two different initial conditions, one with the surfactants randomly distributed (Figure 5a) and another with the surfactants pre-assembled in micelles (Figure 5b). We recall that the surfactant concentration was the same in both cases. The results obtained from both initial conditions are also shown in Figure 5.

**FIG. 5.**
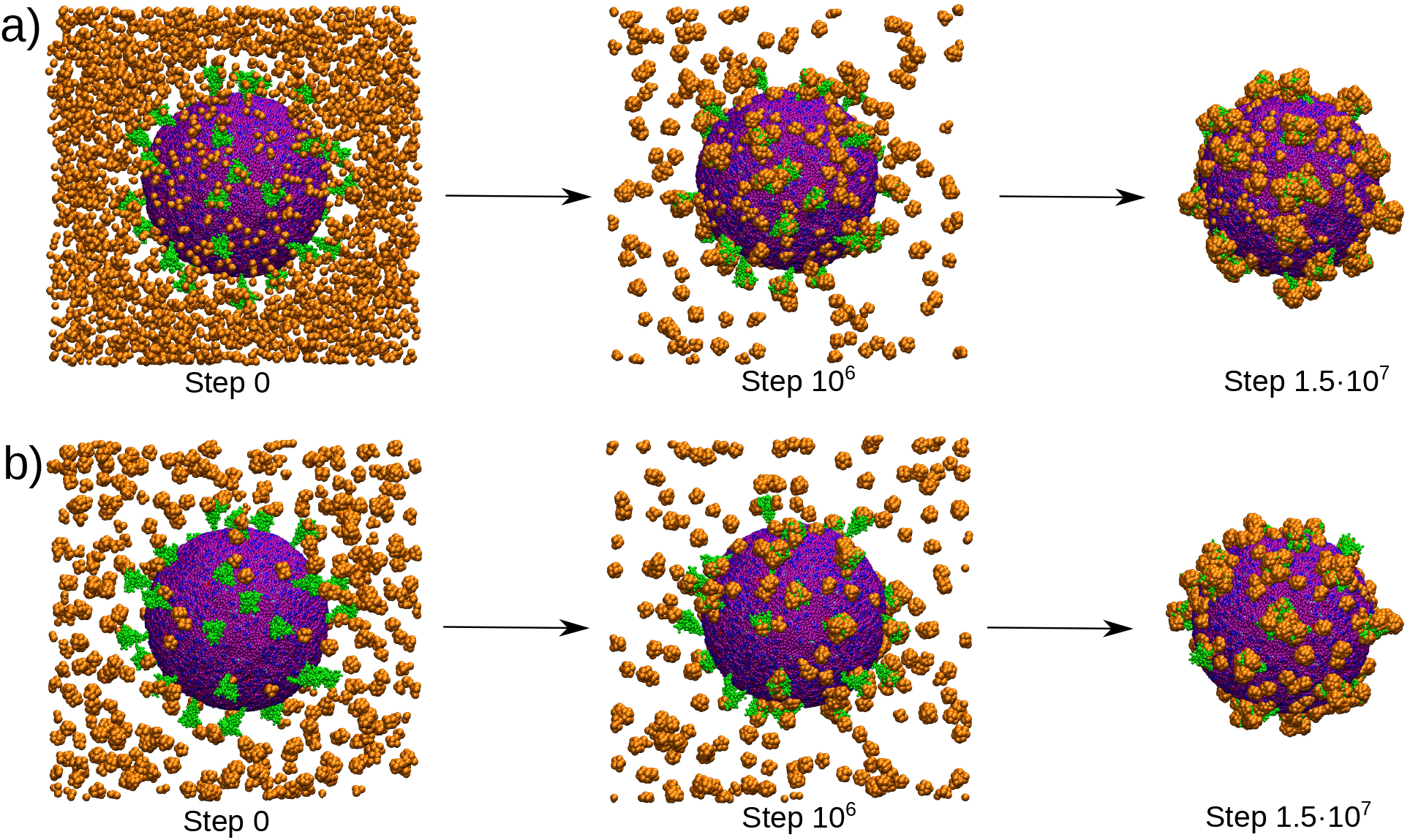
Snapshots of the initial (0 steps), intermediate (after 10^6^ MD steps) and final (1.5 × 10^7^ steps) configurations of Simulation 4 in Table I corresponding to the same simulation with two different initial conditions: a) Initially, the surfactants are randomly dispersed in the medium and b) surfactants are initially pre-assembled into equilibrium micelles in the initial configuration.

In the case of initial random dispersion of surfactants (Figure 5a), the surfactants rapidly form micelles and interact with the virion. At the end of the simulation there are no free surfactants remaining in solution and the integrity of the virus particle was preserved during the simulation (Figure 5a). The same results were obtained for the case of an initial dispersion of surfactants pre-assembled in equilibrium micelles (Figure 5b).

Some interesting features of the simulation can be observed from the snapshots. As seen in Figure 5, after equilibration, the spike proteins are covered by surfactants and also some surfactants are adsorbed onto the envelope membrane. In Figure 6 we show in detail the evolution of the inactivation/coverage of an spike protein by surfactants (see also the movie provided in the supporting information). In 6b we observe how a first micelle gets adsorbed on the spike protein. After some steps (6c), the spike is covered by other micelles, and finally (6d) the spike protein tilts towards the viral envelope probably aided by surfactants that have been previously inserted in the envelope. The process of insertion of surfactants at the envelope is shown in detail in Figure 7 (see also the movie provided in the supporting information). We observe that, after approaching the envelope, a surfactant micelle gets easily adsorbed with the surfactants being incorporated into the membrane, creating a surfactant patch at the membrane exposed to the environment.

**FIG. 6.**
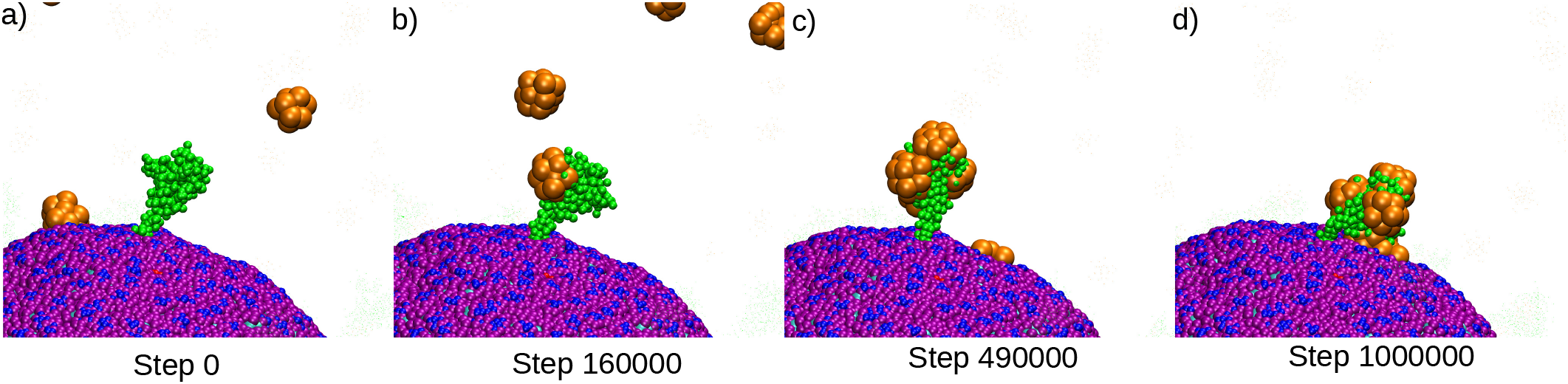
Detail of a snapshot of simulation 4 (see Table I) highlighting the process of coverage by surfactants of a particular virus spike. a) Initial configuration, b) adsorption of a micelle over the spike, c) the spike becomes covered by surfactants, d) the spike modifies its conformation, collapsing over the envelope.

**FIG. 7.**
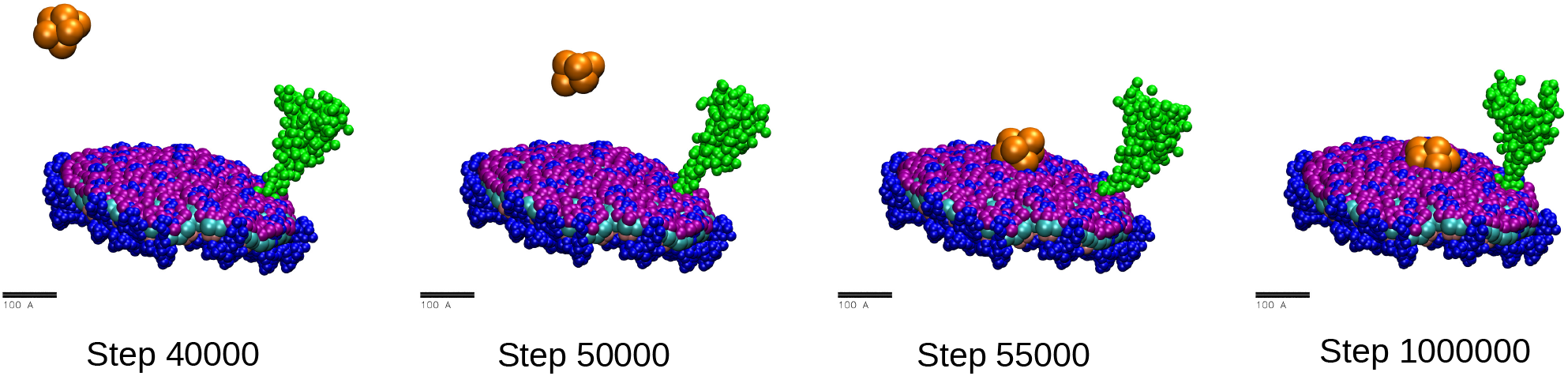
Detail of a snapshot of simulation 4 in Table I highlighting the diffusion of surfactants towards the viral envelope.

A quantitative analysis of the number of surfactants in contact with the S protein and the number of surfactants inserted at the envelope is given in Figure 8. In order to identify the contacts between surfactants and lipids and between surfactants and S proteins, we have first computed their corresponding radial distribution functions. As seen in Figures 8a and 8c, the RDFs show well defined peaks that allow a clear identification of the first coordination shells. We will thus consider that a surfactant is in contact with a lipid or a Spike protein when it is in its first coordination shell. Using these RDFs, we have calculated the number of surfactants in contact with lipids (Figure 8b) and Spike proteins (Figure 8d). As seen in Figure 8c,d, the adsorption of surfactants can be considered equilibrated after 10^7^ time steps. We observe that at equilibrium, the number of surfactants at the envelope (in contact with the lipids) is less than half than the number of surfactants in contact with S protein (≈ 708 and ≈ 1708, respectively). Therefore, in spite of the weakness of the surfactant-S protein interaction, the surfactants are largely adsorbed at the highly exposed S proteins. Also, the simulation reveals that these surfactants not only cover the protein but also affect its orientation at the membrane and its exposure to the environment (see Figure 6). This result indicates the interaction with S as a major mechanism for inaction of the virion particle. This is remarkable, given that in this simulation, the strength of the surfactant-protein attraction is rather weak (of the order of the thermal energy), see Table I. In order to study in detail the impact of the strength of the surfactant-protein and surfactant-lipid interaction, we have considered a full exploration of different values for the parameters in the model (Simulations 1 to 9 in Table I). Since the initial condition had no impact on the final results, we have performed all other simulations in Table I only with the initial condition of randomly distributed surfactants (Figure 5a). The results are summarized in Figure 9 and Table II. In all cases, we obtain adsorption or incorporation of surfactants at the viral particle, which is deformed from its original spherical shape (see snapshots in Figure 9) but maintains its integrity. Overall, the results show a dramatic impact of the interaction between surfactants and proteins, being this interaction more relevant than that between surfactants and lipids. This can be clearly seen in the data shown in Table II. In the case of simulations 1-3 (no interaction between surfactants and proteins), most of the surfactants remain free in solution even for the case with the strongest surfactant - lipid interaction (recall Table I for the strength of the interactions). As long as a some interaction between the surfactants and proteins is included, a massive condensation of surfactants over the virus is observed (even for the cases with a weak surfactant-protein interaction of the order of the thermal energy).

**FIG. 8.**
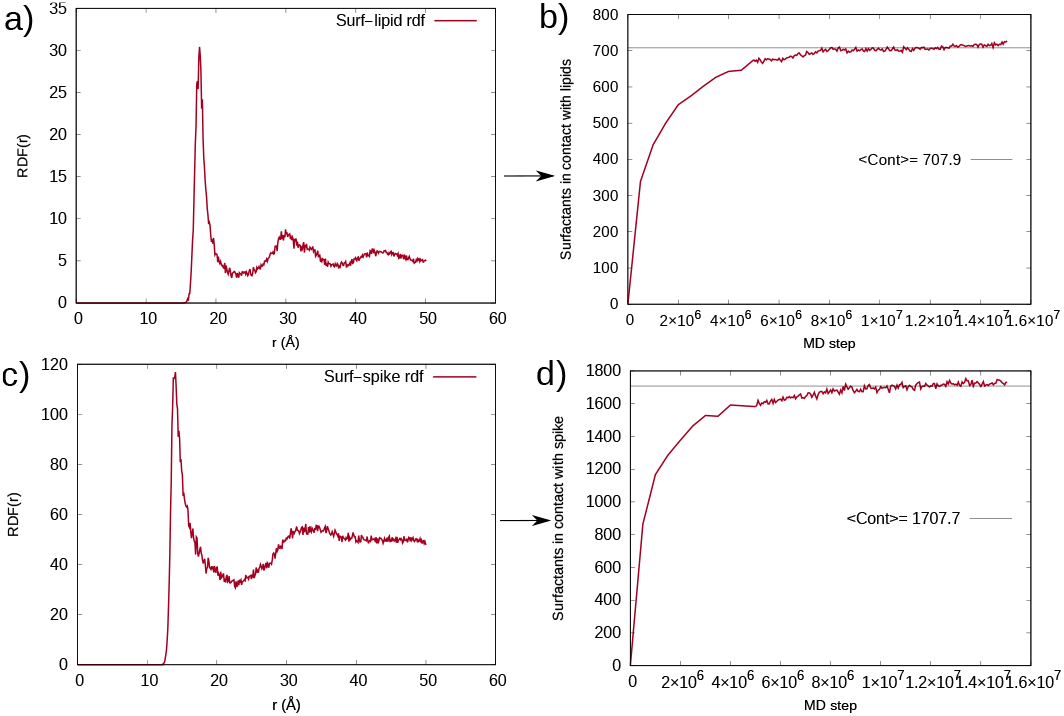
Analysis of Results for Simulation 4 in Table I. (a) Radial distribution function (RDF) between surfactant tails and the hydrophobic beads of the lipids (b) Number of surfactants in contact with lipids (i.e. in the first coordination shell) as a function of time, as identified from the RDF in (a). (c) Radial distribution function (RDF) between surfactant beads and Spike protein beads. (d) Number of surfactants in contact with Spike protein (i.e. in the first coordination shell) as a function of time, as identified from the RDF.

**FIG. 9.**
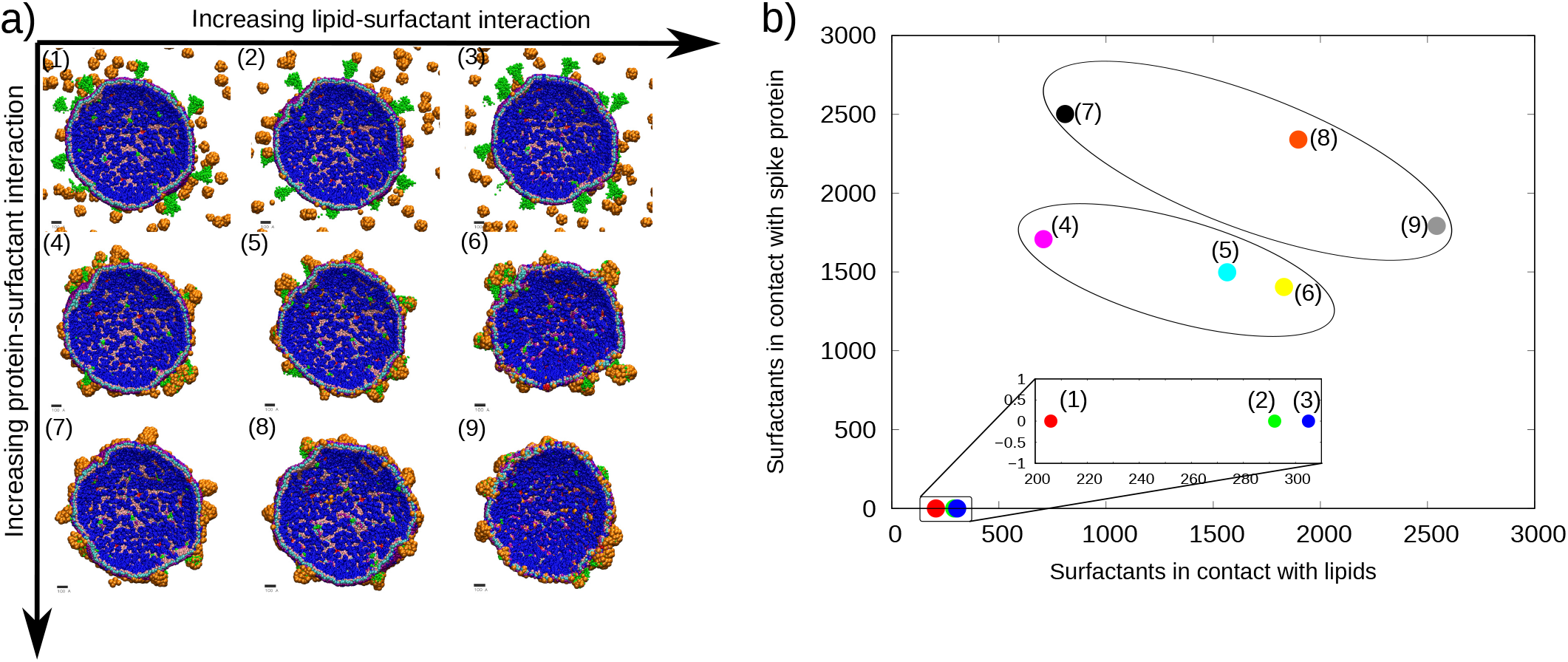
a) Final snapshots of simulations 1 to 9 corresponding to Table I. View of the internal part of the virus in each case. Lipid-surfactant interaction increasing from left to right, and protein-surfactant interaction increasing from top to bottom; b) Number of surfactants in contact with the spike protein vs surfactants in contact with viral lipids for each of the nine simulations.

It is interesting to look in more detail to the number of surfactants that are in contact with lipids and in contact with the Spike protein for each simulation (Table II and Figure 9b).

For simulations 1-3, the contacts between the spike protein and surfactants are nonexistent as there is no attractive interaction between them. The number of surfactants at the envelope increases as the surfactant-lipid interaction increases from simulation 1 to 3, but as we said these values are much lower than in any other simulations including surfactant - Spike interactions.

The results change dramatically when including a weak surfactant - protein interaction, (simulations 4-6, Table I). We have not only a large adsorption of surfactants over the Spike protein but also a substantial increase in the contacts between lipids and surfactants as compared with that obtained in simulations 1-3 (see Table II and Figure 9b). This result indicates that the addition of the surfactant-protein interaction induces a higher interaction between the surfactants and the viral lipids (we recall that the strength of the surfactant -lipid interaction is kept the same in simulations 1 and 4, 2 and 5 and 3 and 6, see Table I). As seen in the snapshots of Figure 9, the virus particles have their spikes completely covered by surfactants, as in the case discussed earlier in this section. The adsorption of surfactants over the S protein is similar for the three simulations but decreases from simulation 4 to 6, as the surfactant-lipid interaction increases. Also, the number of surfactants in contact with the lipids increases from simulation 4 to 6 (Figure 9b).

As we increase the surfactant-protein interaction (simulations 7-9), both the surfactant-Spike and surfactant-lipid contacts increase (Figure 9b). The contacts between surfactants and the spike protein are the highest in simulation 7, which corresponds to the simulation with the highest surfactant-protein interaction combined with the lowest lipid-surfactant interaction.

In addition to Simulations 1-9, we have also performed additional simulations (simulations 7’, 8’ and 9’ in Table I) in order to test the influence of the parameters employed for simulating the supramolecular structure of the SARS-CoV-2 virus in the results. In this additional simulations, we have a weakened lipid-protein interaction (as compared with the original model). In these simulations 7’-9’ we consider that the surfactant-protein interaction is equal to the lipid-protein interaction, as in our previous simulations 7-9 (see Methods for details).

A comparison of the results obtained in simulations 7’-9’ with those obtained in simulations 7-9 is shown in Figure 10. The most strinking difference obtained with the modified (“weakened”) model of virus is that now it is possible to observe a loss of integrity in the virus particle. This is observed in the case of simulation 9’(see Figure 10a) where a spike protein has been detached by the action of surfactants. Compared with the previous simulations, in the case of simulation 9’ we observe a larger deformation of the envelope (Figure 10a). It should be noted that the number of surfactants in contact with the lipids and in contact with the S protein is smaller in simulation 9’ as compared with simulation 9 (Figure 10b). Therefore, a smaller number of adsorbed surfactants produce a larger structural effect in the virus. In the case of simulations 7’ and 8’ the virus envelope retains a nearly spherical shape in contrast with the results in simulations 7 and 8 in which we observe substantial deformation of the envelope (Figure 10a). The number of surfactants in contact with the spike decreases in simulations 7’ and 8’ as compared with 7 and 8, respectively (Figure10b). In fact, the number of surfactants in contact with lipids and proteins obtained in simulations 7’-9’ is more similar to that obtained for simulations 4-6 than for those obtained in simulations 7-9 (see Figures 9b and 10b). This is again in line with the conclusion that the most relevant parameter to interpret our simulations is the value of the surfactant-protein interaction.

**FIG. 10.**
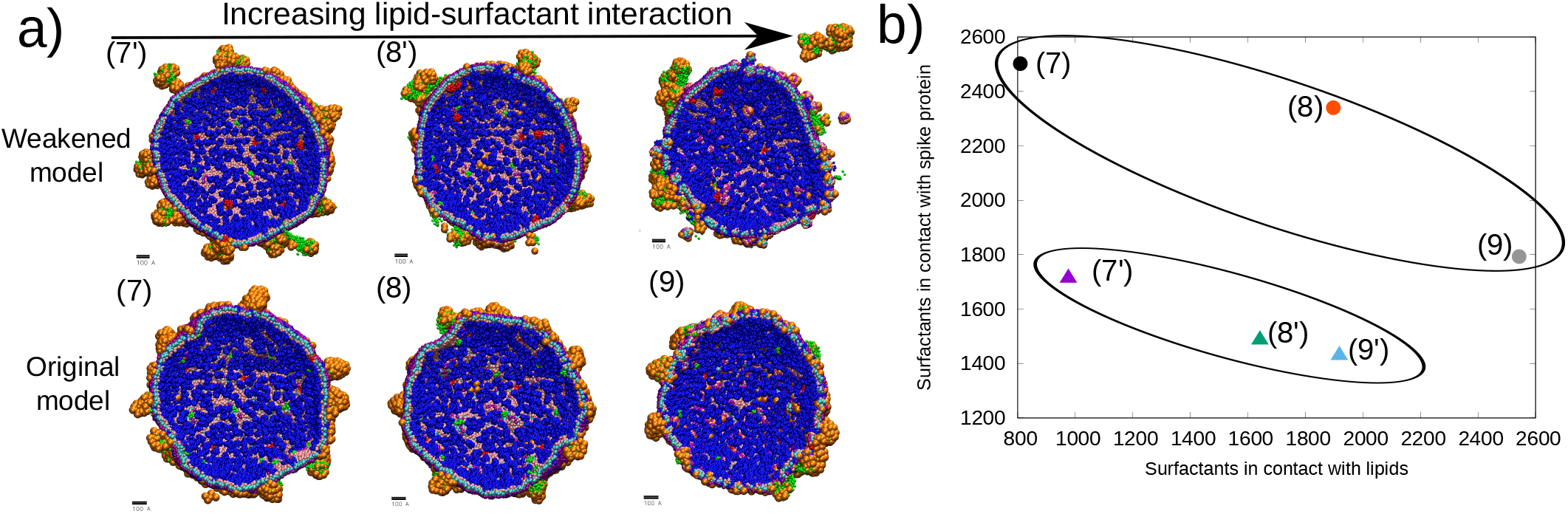
a) Final snapshots of simulations 7’, 8’, 9’, 7, 8, 9 corresponding to Table I. View of the internal part of the virus in each case. Lipid-surfactant interaction increasing from left to right; b) Number of surfactants in contact with the spike protein vs surfactants in contact with viral lipids.

All the simulations discussed so far have been carried out at a constant temperature of 300K and the results were assumed to correspond to final equilibrium configurations. However, it could be possible that these configurations may actually correspond to a metastable state in which the system has become trapped. In order to explore this possibility we have repeated one of the simulations (simulation 7 in the table I) using replica-exchange (RE) technique in order to improve the sampling of configurations. The results from RE simulations are then used as initial condition for subsequent standard simulation at 300K (see Methods for details).

The results of RE simulations are shown in Figure 11. During the RE simulations the 24 different replicas sampled configurations corresponding to temperatures between 300K and 760K (Figure 11b). A final snapshot of the replica ending at 300K is shown in Figure 11a. As in the previous case of nonbiased MD simulations (simulation 7 in Figure 9) we observe a substantial deformation of the shape of the virus with the envelope retaining its integrity without any hole in the structure. The main difference with the previous results is that we see that some viral lipids are removed from the virus envelope (Figure 11a). Concerning the surfactants, some of them are incorporated in the envelope structure, others are adsorbed over the virion and others are still forming micelles in solution (in some cases interacting or encapsulating virion lipids).

**FIG. 11.**
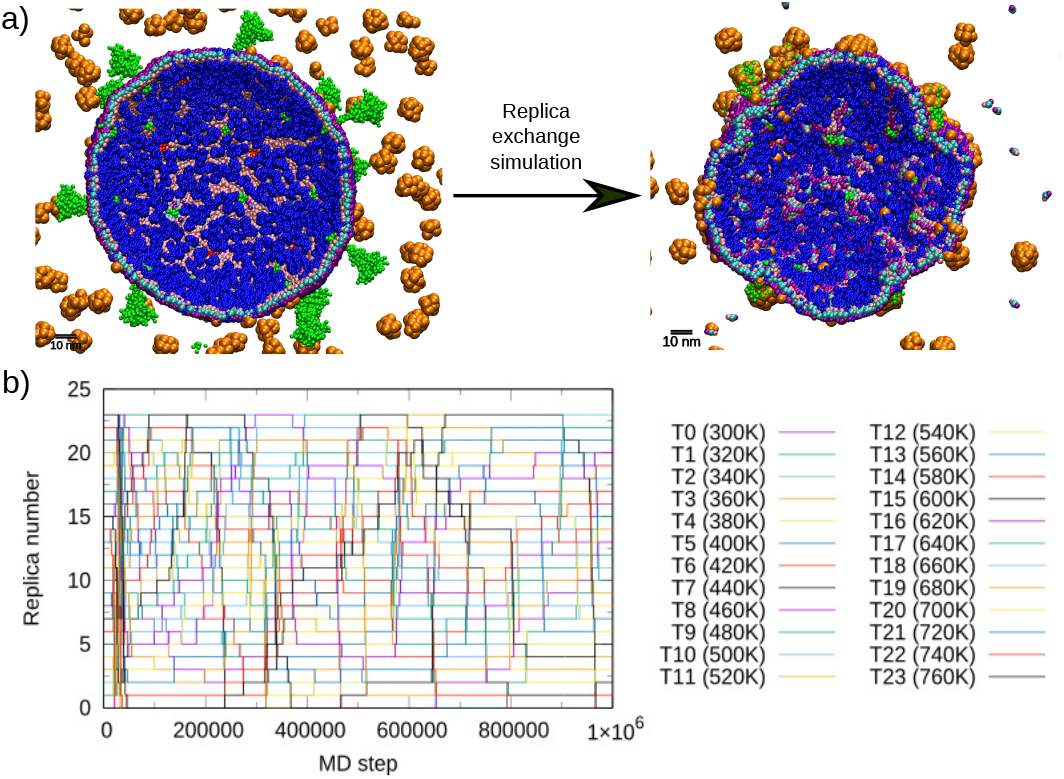
a) Initial and final configuration of the RE simulation. The final configuration corresponds to the replica at 300K at time step 1×10^6^ b) Time evolution of the replica exchange simulation. We show which thermostat corresponds to each system replica as a function of time. Each thermostat is indicated with a different color.

Using the last configuration corresponding to the replica at 300K we run an additional MD simulation run of 1.5 · 10^7^ time steps at 300K. The results are shown in Figure 12. We see that after the MD run the lipids that were removed from the envelope returned from solution to the virus. The incorporation of these lipids to the membrane is not perfect and in the snapshot we see some of them at the surface of the membrane or surrounded by the surfactants. The shape of the envelope remains deformed far from the initial spherical shape. The surfactants that were in solution get adsorbed on the virus. We calculated the number of surfactants in contact with the virion lipids and the spike protein over time as described in the Methods section. At equilibrium, the number of surfactants in contact with virion lipids and the spike protein was ~ 1145 and ~2230, respectively. Comparing this result with that obtained for simulation 7 (Table II), we see that the number of contacts with the lipids has slightly increased and the contacts with the spike have decreased. This is due to the fact that the virial lipids got dissolved during the RE simulation and consequently surfactants could interact better with the lipids, increasing then the number of contacts with the lipids.

**FIG. 12.**
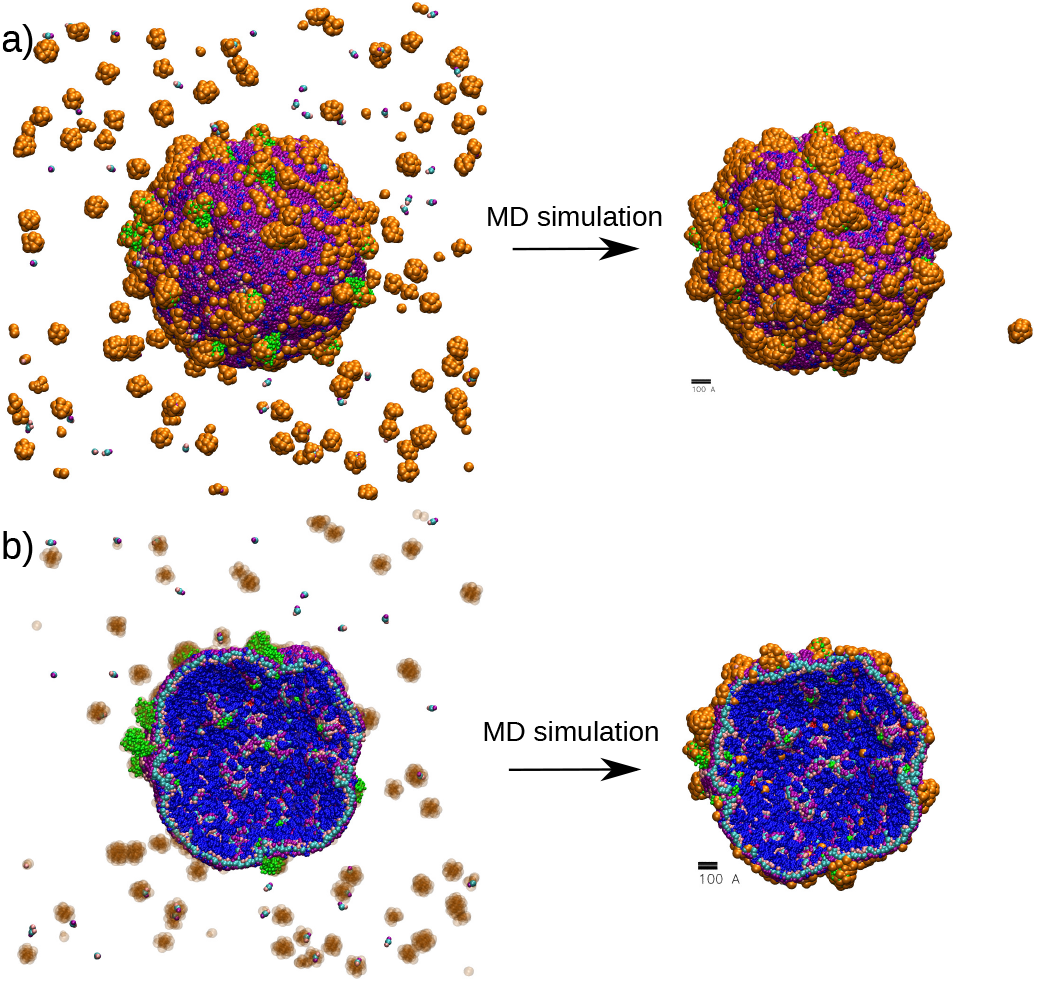
Effect of an unbiased MD simulation at 300K over the configuration obtained from the RE simulation. (a) Full virus and (b) Cuts showing the internal structure of the virus envelope. The images on the left correspond to the last configuration at 300K of the RE simulation. The right images correspond to the last configuration after the MD simulation at 300K.

### B. AA simulations of the interaction of surfactants with SARS-CoV-2 spike protein

Our study of the interactions between the virion and surfactants is completed by doing AA simulations of an inserted spike protein into a membrane and surfactants dissolved in the medium. We consider two types of surfactants, cationic DTAB and anionic SDS (see Methods). The results are shown in Figure 13.

**FIG. 13.**
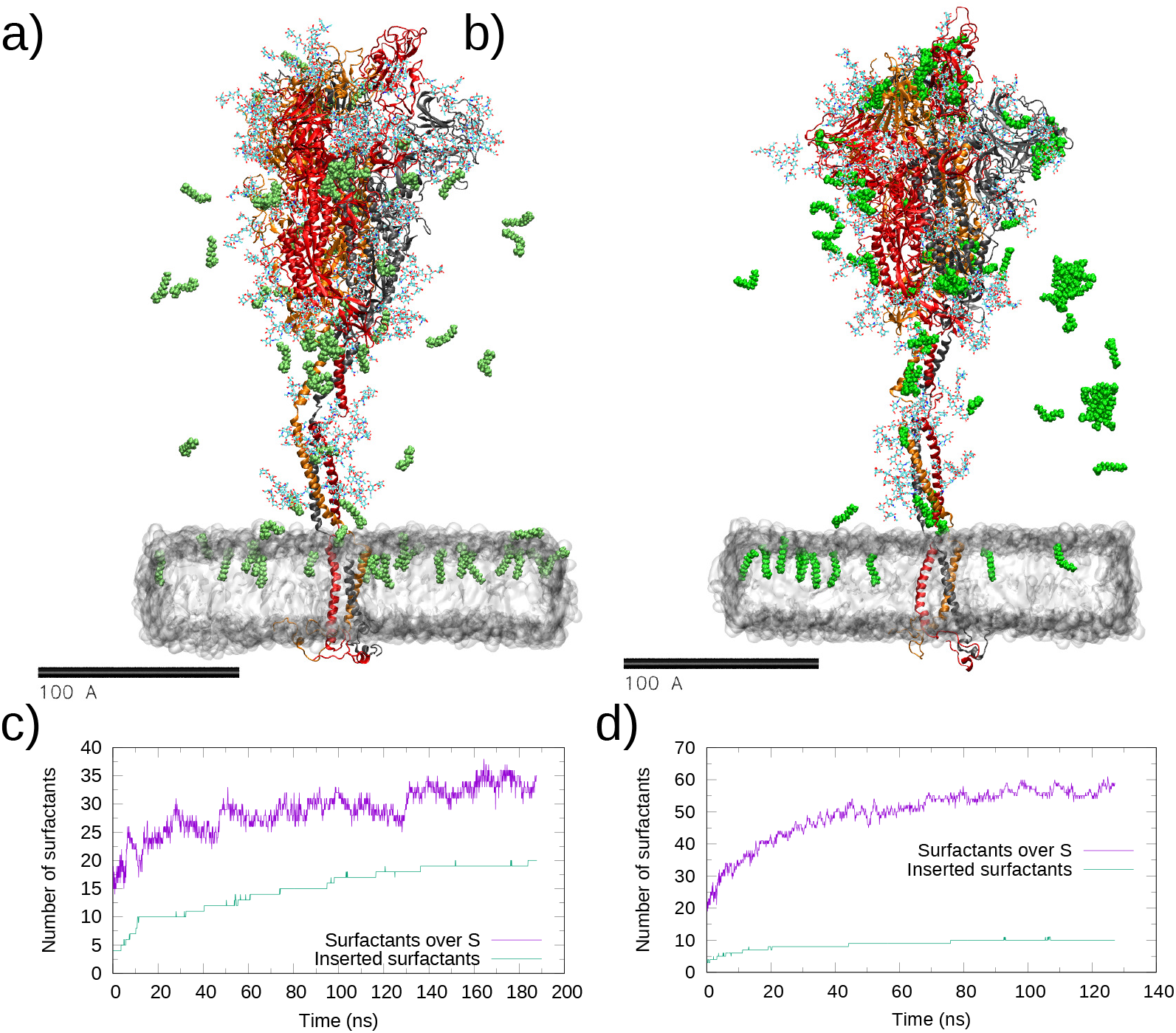
Results from All Atomic simulations (Table III). We show representative snapshots of the simulations with DTAB surfactant (*t* = 169 ns) (panel a) and SDS surfactants (panel b) (*t* = 109 ns). The surfactants are shown in green. For the sake of clarity, water and ions are not shown and we only show the surfactants closer to the virus structure. c) Time evolution of the number of DTAB surfactants in contact with the S protein (purple) and inserted into the membrane (green). d) Time evolution of the number of SDS surfactants in contact with the S protein (purple) and inserted into the membrane (green).

We obtain that both cationic and anionic surfactants interact with the protein and the membrane (Figure 13a,b). It is also interesting to note that for both cases we obtain more surfactants in contact with the S protein than inserted into the lipid membrane (Figure 13c and 13d). In the case of DTAB surfactants (Figure 13c), we obtain equilibrium values of ~ 20 surfactants inserted into the membrane and ~ 35 surfactants adsorbed at the spike protein. In the case of SDS surfactants (13d), we obtain ~10 surfactants inserted into the membrane and ~ 55 surfactants adsorbed over the Spike protein.

The fact that we have more DTAB than SDS surfactants inserted into the membrane could be due to the fact that the membrane is formed by neutral and negatively charged phospholipids.

More interesting is the fact that the number of SDS surfactants interacting with the S protein is much higher than those of DTAB, in spite of the overall negative charge of the spike protein. In this case, SDS surfactants, also adsorb at the Receptor Binding Domain (RBD) of the spike protein, meaning that this type of surfactant may inactivate the infectivity of the virion by attacking this spot. However, in the case of DTAB we did not observe surfactants adsorbing at the RBD. In Figure 14 we show a zoom of the SDS surfactants over the RBD domain of the spike protein. Seven SDS surfactants are directly interacting with the RBD monomer in “up” conformation. In the second image we see which specific residues of the RBD interact with one of the surfactants. The surfactant interacts with Spike hydrophobic side chains (LEU455, PHE490, ILE46, TYR421). Interestingly, the positively charged NH_3_ group in LYS417 interacts with the negatively charged head of a second SDS surfactant, as seen in the snapshot. In this model of S protein, the net charge of the RBD is neutral, so the hydrophobic and local electrostatic environment should play a role for any binding site, as suggested in^19^.

**FIG. 14.**
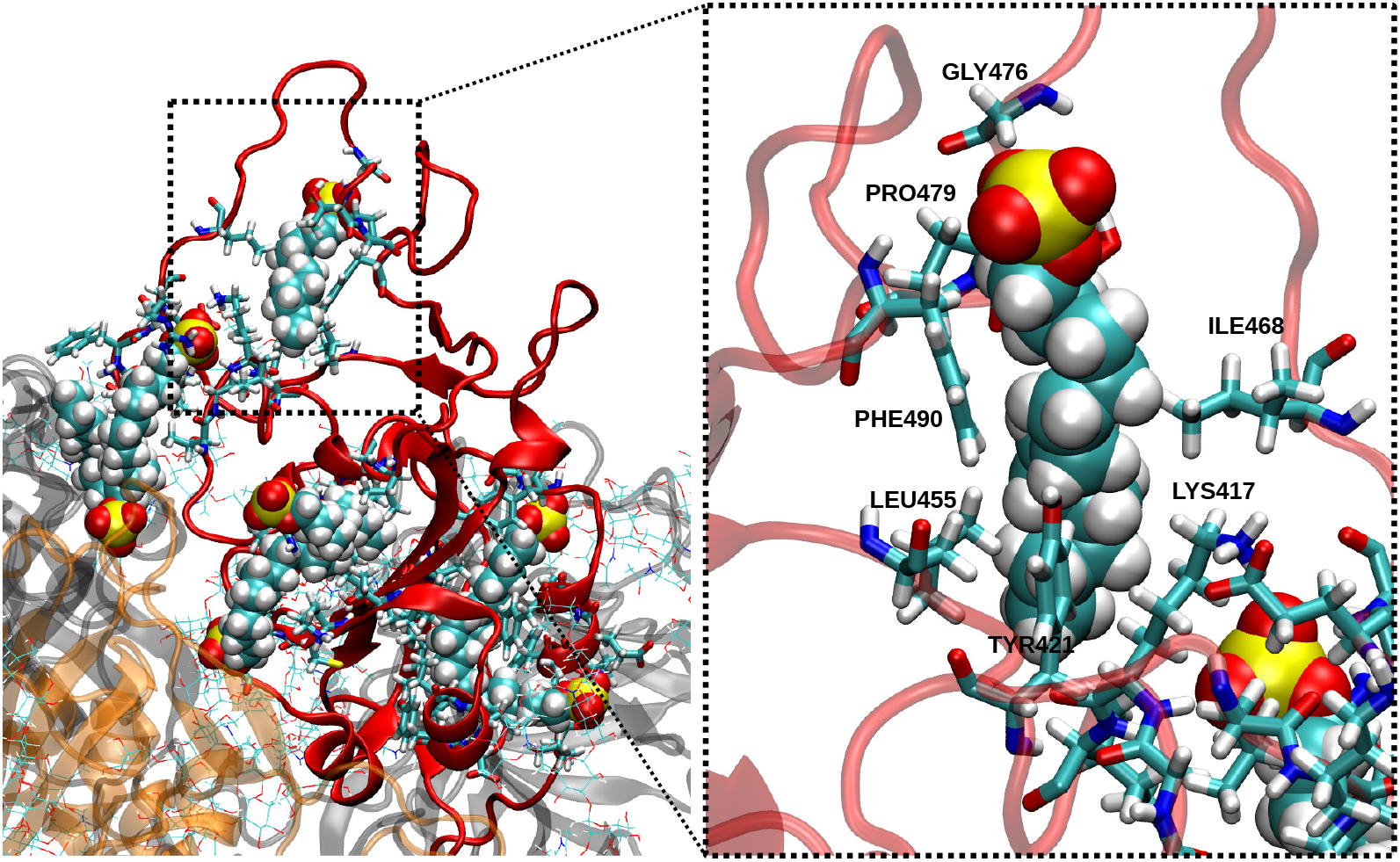
Representative snapshot (t = 109 ns) of the AA simulation of the S protein with SDS surfactants showing the detail of the interaction of SDS surfactants with the spike protein RBD. The S monomer in the “up” conformation is shown in red using the cartoon representation. The emphasized region highlights the interaction of one SDS surfactant with the S residues. protein residues are shown in licorice representation, surfactants as Van der Waals spheres and glycans as lines (color codes: carbon in cyan, hydrogen in white, nitrogen in blue, oxygen in red, sulfur in yellow).

The results obtained for the adsorption of the SDS surfactant over the Spike protein are similar to those obtained in Simulation 4 of the CG model (Table II. In that case, the number of surfactants interacting with the spike protein was 1708, which means there are ~57 surfactants in contact with each spike protein. Even though the differences between CG and AA simulations, we see that the order of magnitude of the number interacting surfactants with the S protein is the same.

## IV. CONCLUSIONS

In this work, we presented CG molecular dynamics simulations of the interaction between surfactants and an entire SARS-CoV-2 virus. We also performed AA simulations between surfactants and the spike glycoprotein inserted into a membrane patch.

The main result obtained from the CG simulations is that the most important effect of surfactants over the virus is to block the spike proteins, thus cancelling their function. Even if the strength of the interaction between the surfactants and the S protein is weak (of the order of the thermal energy), surfactants cover abundantly the spike proteins and make them collapse onto the envelope surface. This collapsing mechanism was observed to be aided by surfactants that were previously attached at the envelope. Moreover, in most of CG simulations, we observed a substantial deformation of the virus envelope from its spherical shape, but in none of them a pore formation or a rupture of it was produced. We conclude that the interaction between the surfactants and the spike proteins is even more crucial than the interaction with viral lipids when we proceed to the virus inactivation by means of surfactants.

Concerning the all atomic simulations, we observed that both for anionic and cationic surfactants, the number of surfactants in contact with the spike protein was higher than the number of them inserted into the membrane, which is consistent with the results of the CG simulations. Interestingly, anionic surfactants have a higher affinity with the spike protein, specially with the RBD domain. This fact implies that the choice of a specific surfactant is meaningful when we want to interfer on the infectivity of the virus by attacking the spike protein.

Some similarities can be seen when comparing the CG and AA simulations. Surfactants interact with both the S protein and the membrane, inferring that the mechanism of inactivation of SARS-CoV-2 virus by surfactant means is probably not unique. AA simulations support the idea that the S protein is a key target for surfactants and that the high exposure of the protein in the environment captures the surfactants on the surroundings. Actually, we observed how in the case of SDS surfactants, they interact with the S protein with hydrophobic and electrostatic interactions. However, in AA simulations we cannot observe the effect on the S protein seen in the CG simulations (the tilt of the S protein on the virion envelope). Due to the large size of the simulated system it is certainly hard to observe this type of behaviour in AA models.

At this point, it is also interesting to propose possible experimental verification of the predictions presented here. Experimentally, it seems very difficult to be able to observe directly (by suitable microscopy techniques such as cryo-TEM) the blocking of the S protein predicted by our simulations. But biophysical studies of suspensions of the full S protein or the RBD of S in presence of anionic and cationic surfactants seem feasible, and these may reveal the different binding affinity of these surfactants suggested by our simulations.

Of course, it is important to keep in mind the limitations of the used computational models and simulations. Regarding the CG simulations, the most important limitation is the loss of resolution with respect to an atomistic model since a single CG bead in this model corresponds to tens of atoms. Future work could be directed to improve the link between CG and atomistic simulations, for example deriving the interaction parameters of the CG model of the surfactants from atomistic simulations. And therefore, it makes us miss details that can happen at the molecular level. In the AA simulations, we considered the spike inserted into a lipid bilayer but the envelope contains other proteins (M and E proteins) can also be targets of surfactants. Ongoing work on the study of the effect of surfactants on models that also consider the M proteins of the envelope is now being carried out.

As a final remark, we would like to point out an interesting possibility for high throughput screening of surfactants effective against SARS-CoV-2 that arises from the present results. On one hand, our simulations suggest as the best strategy for the design of surfactants as virucidal agents to focus on those strongly interacting with the S protein. On the other hand, a recent study^38^ shows that computational docking techniques can be successfully employed for the screening of large databases of compounds identifying their possible interactions with the S protein of SARS-CoV-2. The combination of these two results suggest as a strategy for selecting surfactants for disinfecting products the use of docking techniques for the fast screening of databases of surfactants to identify those with stronger interactions with S, in a way analogous to that employed in Ref^38^.

## ACKNOWLEDGMENTS

This work was supported by the Spanish Ministry of Science and Innovation through Grant No. PID2021-124297NB-C33, the “Severo Ochoa” Grant No. CEX2019-000917-S for Centres of Excellence in R&D awarded to ICMAB and the FPI grant PRE2020-093689 awarded to M. D. We thank the CESGA supercomputing center for computer time and technical support at the Finisterrae supercomputer.

We thank Dr. Alvin Yu and Prof. G.A.Voth (University of Chicago) for their help on the implementation of the virus model and for providing us the code for the “Two Gaussian” potential.

M. D. is enrolled in the Material Sciences PhD program of the Universitat Autonoma de Barcelona.

## AUTHOR DECLARATIONS

### Author Contributions

M.D. and J.F. designed the study and simulations. M.D. performed the simulations and analysed the data supervised by J.F. M.D. made the Figures. M.D and J.F. discussed the results and wrote the manuscript.

## DATA AVAILABILITY STATEMENT

Input files for the LAMMPS CG simulations, the parameters of the CG force field and coordinate and structure files for our all atomic NAMD simulations can be found at our GitHub repository at [GitHub ICMAB SoftMatter](https://github.com/soft-matter-theory-at-icmab-csic). We also provide sample scripts of the simulations.

## Appendix: Implementation of the Soft Grime potential

The equations employed for the force field described in the Methods section were implemented in the standard release of LAMMPS, except the two Gauss potential and the soft potential for lipids and surfactants introduced by^25^ (Eq.(4). This potential has to be introduced by adding an additional code, available at^27^. In order to ensure reproducibility of our results, we discuss here the equations implemented in the code, since most of that information cannot be found in the bibliography.

The implementation of this interaction is based on the following equation for the force between two beads:

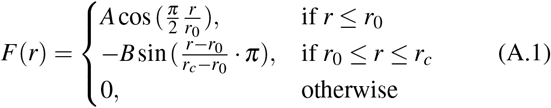

This interaction is repulsive for *r* < *r*_0_, attractive for *r*_0_ < *r* < *r_c_* and is switched off for *r* > *r_c_*. The value of *r*_0_ is taken as the size of the bead and *r_c_* is taken as *r_c_* = 2*r*_0_. The force has two parameters *A* and *B* that determine the strength of the repulsive and attractive force. The potential corresponding to Eq.(A.1) is given by:

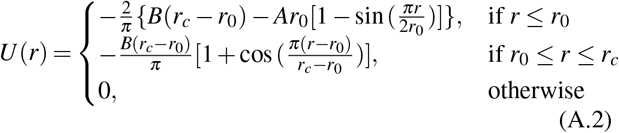

Eqs.(A.1) and A.2 as implemented in LAMMPS require the user to specify the values of *A* and *B*. The relation between the energy parameters *U*_0_ and *U_c_* in Eq.4 as introduced in the Methods section and the *A* and *B* parameters in Eq.A.2 is:

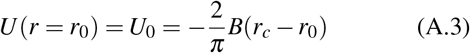

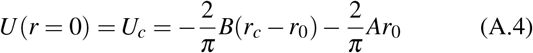

The values of *A* and *B* employed in the simulations of Table I are summarized in Table IV for the sake of reproducibility. For a justification of the selected values, please refer to the Methods section.

**TABLE IV.**
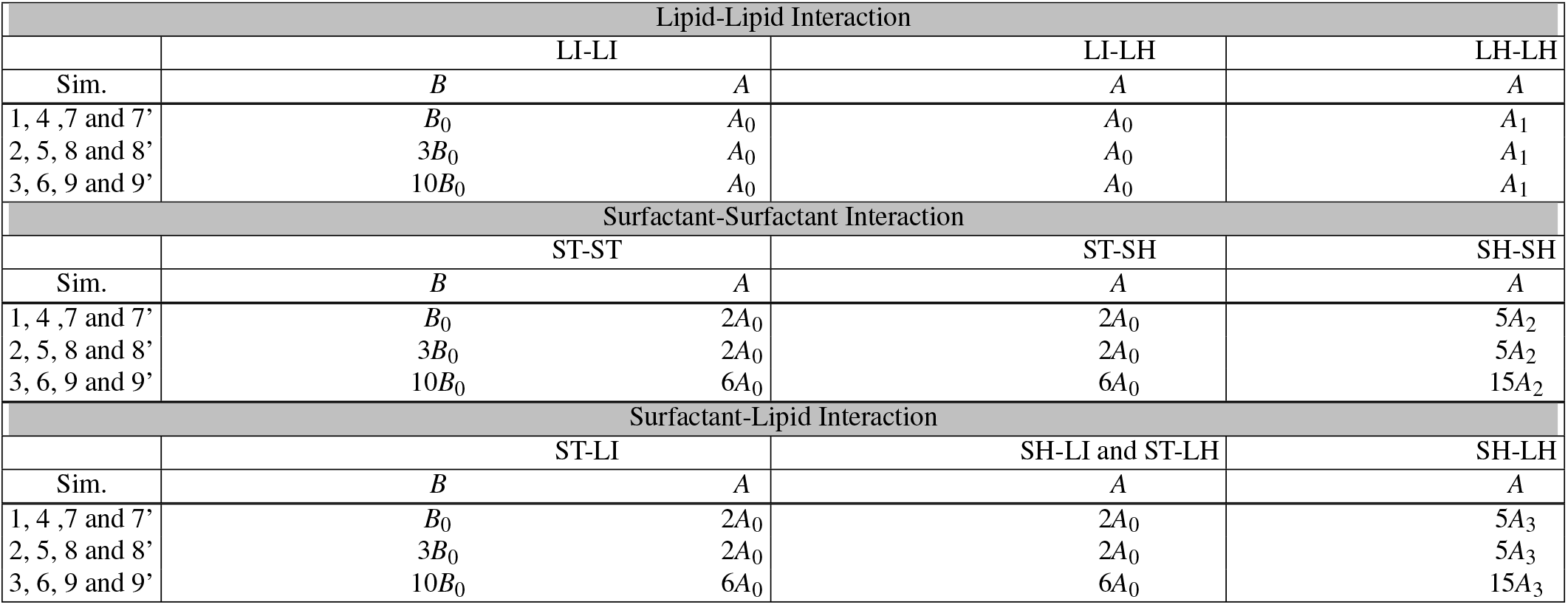
Values of *A* and *B* parameters in Eq.A.2 for the interactions between lipids and surfactant beads used in simulations in Table I. We employ the following abbreviations for the beads: ST = surfactant tail bead, SH = surfactant head bead, LI = lipid interface bead and LH=lipid head bead. We also define the following characteristic values: 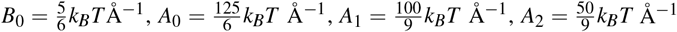 and 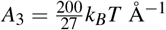.

| Lipid-Lipid Interaction |  |  |  |  |
| --- | --- | --- | --- | --- |
|  | LI-LI |  | LI-LH | LH-LH |
| Sim. | $B$ | $A$ | $A$ | $A$ |
| 1, 4, 7 and 7' | $B_0$ | $A_0$ | $A_0$ | $A_1$ |
| 2, 5, 8 and 8' | $3B_0$ | $A_0$ | $A_0$ | $A_1$ |
| 3, 6, 9 and 9' | $10B_0$ | $A_0$ | $A_0$ | $A_1$ |
| Surfactant-Surfactant Interaction |  |  |  |  |
|  | ST-ST |  | ST-SH | SH-SH |
| Sim. | $B$ | $A$ | $A$ | $A$ |
| 1, 4, 7 and 7' | $B_0$ | $2A_0$ | $2A_0$ | $5A_2$ |
| 2, 5, 8 and 8' | $3B_0$ | $2A_0$ | $2A_0$ | $5A_2$ |
| 3, 6, 9 and 9' | $10B_0$ | $6A_0$ | $6A_0$ | $15A_2$ |
| Surfactant-Lipid Interaction |  |  |  |  |
|  | ST-LI |  | SH-LI and ST-LH | SH-LH |
| Sim. | $B$ | $A$ | $A$ | $A$ |
| 1, 4, 7 and 7' | $B_0$ | $2A_0$ | $2A_0$ | $5A_3$ |
| 2, 5, 8 and 8' | $3B_0$ | $2A_0$ | $2A_0$ | $5A_3$ |
| 3, 6, 9 and 9' | $10B_0$ | $6A_0$ | $6A_0$ | $15A_3$ |

